# Declaration of Fermentation: Community-Embedded Wild Yeast Bioprospecting as a Model for Place-Based CURE Design

**DOI:** 10.64898/2026.06.29.735295

**Authors:** Spencer J. Gray, Kaitlyn Taylor, Kennadi A. Shumaker, Matthew L. Bochman

**Affiliations:** Molecular & Cellular Biochemistry Department, Indiana University, Bloomington, IN 47405, USA

**Keywords:** course-based undergraduate research experience (CURE), microbiology education, wild yeast, *Saccharomyces cerevisiae*, bioprospecting, fermentation, place-based education, community partnership, semi-quincentennial, sensory evaluation

## Abstract

Course-based undergraduate research experiences (CUREs) are widely recognized as a high-impact practice in biology education, yet most existing CURE frameworks treat the research organism as an interchangeable teaching prop rather than a genuine scientific contribution. We argue that place-based, community-embedded CUREs – in which students isolate, characterize, and publicly deploy a locally meaningful wild organism – constitute a qualitatively distinct model warranting broader adoption. As proof of concept, we present the Declaration of Fermentation project at Indiana University Bloomington: graduate researchers isolated a wild *Saccharomyces cerevisiae* strain from the bark of a campus landmark tree, confirmed its wild provenance by whole-genome sequencing and phylogenomics, and partnered with local craft breweries to produce a colonial-era inspired ale released publicly for the 250th anniversary of the Declaration of Independence. Volunteer sensory panels at two independent public tasting events (combined n = 33–34 per attribute) confirmed a fruity-funky profile consistent with wild-strain fermentation, with no significant differences between events (Mann–Whitney U, Benjamini– Hochberg-corrected p > 0.05 for all 11 attributes). We describe three design principles – genomically confirmed strain identity, mandatory community partnership, and place-based historical narrative – that distinguish this model from prior wild yeast brewing CUREs, discuss how these principles generalize to other institutions and fermentation vehicles, and identify next steps for formal learning assessment. Complete implementation protocols are provided as supplemental Appendices 1–6, and the bioinformatics pipeline is freely available at https://doi.org/10.5281/zenodo.20679384.

## INTRODUCTION

Course-based undergraduate research experiences (CUREs) have emerged as a high-impact practice for improving student engagement, retention, and scientific identity in the biological sciences (1, 2). Unlike traditional “cookbook” laboratory exercises, CUREs engage students in genuine inquiry: they generate data that are not already known, navigate experimental uncertainty, and revise their approaches in response to results. These features distinguish CUREs from inquiry-guided laboratories, in which the outcome is predetermined, even if the path is open-ended. Despite widespread recognition of their value, many undergraduate life sciences laboratory courses continue to rely on well-characterized model strains and tightly scripted protocols, limiting opportunities for authentic discovery and reducing student ownership of the research process.

Wild yeast bioprospecting offers a particularly compelling entry point for CURE design. Microorganisms are ubiquitous, diverse, and readily sampled with minimal equipment, yet even simple enrichment strategies can yield strains with novel physiological and metabolic properties (3-6). In the context of fermentation, industrial and research applications have long centered on a narrow set of domesticated *Saccharomyces cerevisiae* strains, but environmental surveys have revealed that wild isolates harbor substantial genetic and functional diversity, including variation in fermentation rate, ethanol tolerance, flavor compound production, and thermal stability (5-9). This diversity translates directly into student-driven research questions: Which isolate ferments most vigorously? Which produces distinct sensory characteristics? Do isolates from different microhabitats differ in their fermentation profiles? Because the answers are not predetermined, students engage with the genuine uncertainty that defines scientific practice.

Two prior publications have demonstrated the feasibility of wild yeast brewing CUREs at the undergraduate level. Scholes *et al*. (2021) describe a guided-inquiry laboratory course organized around isolation, phenotypic characterization, and head-to-head fermentation comparison of wild and commercial *S. cerevisiae* strains, with pre- and post-assessment showing significant learning gains (10). DeHaven *et al*. (2022) report a semester-long CURE integrating standard microbiology protocols – pure culture isolation, staining, microscopy, PCR, and gel electrophoresis – into an authentic brewing project assessed for knowledge gains (11). Both frameworks are valuable, broadly applicable, and appropriately generic: the wild yeast comes from wherever, the beer is brewed with whichever strain performs best, and the educational value lies in the process.

Our project differs in three ways that we argue constitute a qualitatively distinct category of CURE. First, the strain itself is a citable, genome-sequenced, depositable scientific contribution – not a teaching prop – and its wild provenance is verified by whole-genome sequencing rather than assumed from the sampling context. Second, the community partnership with local craft breweries is a structural feature of the curriculum, not an optional extension: students understand from the outset that the product of their research will be experienced by the public, which changes the stakes and meaning of the work. Third, the historical narrative – connecting the isolation of a wild yeast from the “Celebri-tree,” a landmark burr oak on the IUB campus, to the brewing of a colonial-era ale for the 250th anniversary of the Declaration of Independence – provides motivational scaffolding that a generic “yeast from wherever” framework cannot replicate. Place-based learning theory predicts that locally meaningful contexts increase student engagement and knowledge retention (12); our implementation tests whether that advantage can be designed into a CURE from the start rather than added as narrative decoration.

Here, we describe the Declaration of Fermentation project as a proof-of-concept for this model. We present our research question, methods, and results in sufficient detail to support replication and formal assessment at other institutions. We also provide, as supplemental material, the protocols, assessment instruments, and course schedule needed to implement an analogous CURE elsewhere. Our central claim is not that our specific strain or our specific historical event is required. Rather, it is that the design principles of strain-specific identity, community partnership, and place-based narrative are generalizable features that any institution could implement with a locally or historically meaningful equivalent.

## PROOF-OF-CONCEPT IMPLEMENTATION

### The Declaration of Fermentation Project

The Declaration of Fermentation was piloted by three graduate researchers (S.J.G., K.T., K.A.S.) under faculty supervision (M.L.B.) during the 2025–2026 academic year at Indiana University-Bloomington (IUB). The central research question was: can a community-embedded, historically contextualized wild yeast bioprospecting CURE yield a scientifically validated environmental isolate capable of producing a fermentation product with sensory characteristics consistent with wild-strain origin? The project builds on a decade of undergraduate bioprospecting in the Bochman laboratory (5, 6, 9) and represents the first formal integration of those activities into a structured CURE framework with explicit learning objectives, modular assessment scaffolding, and a mandatory community partnership. Full protocols, student handouts, instructor notes, and assessment rubrics are provided in Appendices 1–6; the bioinformatics pipeline is documented at https://doi.org/10.5281/zenodo.20679384 (13).

Environmental samples were collected from multiple IUB campus sites, with priority sampling of a >200-year-old burr oak (the “Celebri-tree”), and enriched for ethanol-tolerant yeasts using YPD8E5 selective medium. Isolates were screened for fermentation performance and identified by PCR amplification and Sanger sequencing of the rDNA D1/D2 domain (14). One isolate, designated *S. cerevisiae* DoF1, was selected for brewing based on robust wort fermentation and confirmed as a genuine wild isolate by whole-genome sequencing (Oxford Nanopore Technology; Plasmidsaurus) and phylogenomic analysis (832-gene supermatrix; IQ-TREE3; LG+G4; 1,000 ultrafast bootstrap replicates) (15, 16). DoF1 was transferred to Upland Brewing Company (Bloomington, IN) for commercial-scale fermentation; the finished ale was distributed publicly by five local craft breweries in June 2026. A volunteer sensory panel evaluated the beer using a structured scorecard (Appendix 5) at a public tasting event; all work was conducted at BSL-1 and the sensory evaluation was determined by the IU Office for Research Compliance not to constitute human subjects research.

### Curriculum Structure

The CURE is organized into six modules spanning one to two semesters (Table 1): 1) historical and contextual framing; 2) environmental sampling and enrichment; 3a) molecular identification; 3b) whole-genome sequencing and phylogenomics (advanced extension); 4) small-scale fermentation and sensory evaluation; 5) commercial brewery partnership and public tasting event; and 6) capstone data synthesis and communication. The framework is modular – resource-limited settings can omit Module 3b or Module 5 without loss of core CURE features – and maps naturally onto an interdisciplinary implementation involving students from history, field ecology, food science, bioinformatics, and communications (Appendix 4). Intended for upper-division undergraduates in microbiology, biochemistry, or related programs, the core CURE requires only standard microbiological equipment plus access to an external Sanger sequencing service. Prerequisite knowledge includes aseptic technique, PCR, and gel electrophoresis; no bioinformatics experience is required for Module 3b.

**Table 1.** Declaration of Fermentation CURE: module overview. Module 3b (whole-genome sequencing and phylogenomics) and Module 5 (full-scale brewery production) are optional extensions for advanced courses or up to two-semester implementations. Week ranges are approximate; some modules can run in parallel.

| Module | Phase / Weeks | Activities | Key Skills | Assessment |
| --- | --- | --- | --- | --- |
| 1 | Pre-lab<br>Weeks 1–2 | Historical and contextual framing; recipe sourcing and translation from primary sources; site selection for environmental sampling; sampling permit logistics; safety training | Primary source research; scientific writing; field safety; microbial ecology concepts | Site selection rationale (written); annotated historical recipe with modernization notes |
| 2 | Weeks 2–4 | Environmental sampling (bark swabs, soil, fruit); YPD8E5 enrichment medium preparation; inoculation and incubation; colony isolation on WLN agar; colony morphology documentation; Gram stain and microscopy | Aseptic technique; enrichment culture; streak-plate isolation; brightfield microscopy; laboratory notebook keeping | Laboratory notebook entries; colony morphology data table; microscopy images with annotations |
| 3a | Weeks 4–6<br>(core track) | Genomic DNA extraction; PCR amplification of rDNA D1/D2 domain; gel electrophoresis; Sanger sequencing submission and result interpretation; BLASTN species identification | DNA extraction; PCR design and troubleshooting; gel electrophoresis; sequence analysis; database searching | Annotated gel image; BLASTN results with interpretation; species identification report |
| 3b | Weeks 4–6<br>(advanced extension) | ONT whole-genome sequencing submission; BUSCO assembly assessment; AUGUSTUS gene annotation; MAFFT alignment; trimAl trimming; IQ-TREE3 phylogenomic analysis; interpretation of phylogenetic placement relative to wild and domesticated reference strains | Genome assembly quality assessment; gene prediction; multiple sequence alignment; maximum likelihood phylogenetics; command-line bioinformatics tools | BUSCO completeness report; annotated phylogenetic tree with written interpretation; peer-reviewed draft methods paragraph |
| 4 | Weeks 6–12<br>(fermentation) | Small-scale wort fermentation (2 weeks); hydrometer/refractometer gravity monitoring; pH measurement; sensory evaluation of green beer; bottle conditioning in 12-oz glass bottles (2 weeks); final structured sensory evaluation using scorecard instrument | Fermentation monitoring; data collection and graphing; sensory evaluation methodology; statistical summary (mean ± SD) | Fermentation data table and graph (attenuation over time); completed sensory scorecards; written fermentation report |
| 5 | Weeks 8–16<br>(full-scale;<br>optional) | Brewery partnership<br>coordination; recipe finalization<br>with brewer; commercial-scale<br>fermentation; quality control<br>sampling; public tasting event<br>logistics; structured volunteer<br>sensory panel; community<br>outreach | Professional communication;<br>project management; sensory<br>panel design; community<br>science engagement; data<br>aggregation and presentation | Project management log;<br>brewery partnership<br>reflection; sensory panel<br>dataset; public-facing<br>summary poster or social<br>media content |
| 6 | Final weeks<br>(capstone) | Data synthesis across all<br>modules; scientific poster or<br>oral presentation; peer review of<br>written reports; reflection on<br>research process and<br>community impact | Scientific communication;<br>peer review; reflection and<br>metacognition; data synthesis<br>across disciplines | Poster or oral presentation;<br>peer review completed;<br>individual reflection essay;<br>pre/post CURE assessment<br>instrument |

## KEY FINDINGS

### Genomic Validation of Wild Provenance

The DoF1 assembly comprised 20 contigs spanning 11.92 Mb at 183× mean coverage depth (contig N50: 795 kb). BUSCO completeness against saccharomycetes_odb10 was 99.4% (97.4% single-copy, 2.0% duplicated, 0.6% missing) (17), indicating a near-complete diploid nuclear genome. Gene annotation identified 5,408 predicted protein-coding genes with best BLAST hits predominantly to *S. cerevisiae* S288c (96.5%), with secondary hits to environmental, vineyard, and clinical isolates rather than laboratory strains. The *URA3* locus appears intact and flocculation genes (*FLO8, FLO9*, and *FLO10*) are present, both of which are features absent or disrupted in domesticated laboratory strains, further supporting wild provenance. Phylogenomic analysis placed DoF1 within a well-supported clade (100% bootstrap) comprising wild North American strains (YPS163, YPS1009) and wine-lineage strains (AWRI1631, CLIB324), entirely separate from laboratory, bioethanol, and West African lineages (Fig. 1). *S. cerevisiae* is not considered a food safety risk in healthy individuals (18), and this placement is fully consistent with use in a public fermentation product. The genome has been deposited in NCBI GenBank and the SRA under BioProject PRJNA1477971 (BioSample: SAMN60825368; GenBank: JBZFQT010000000; SRA: SRR39135318).

**Figure 1.**
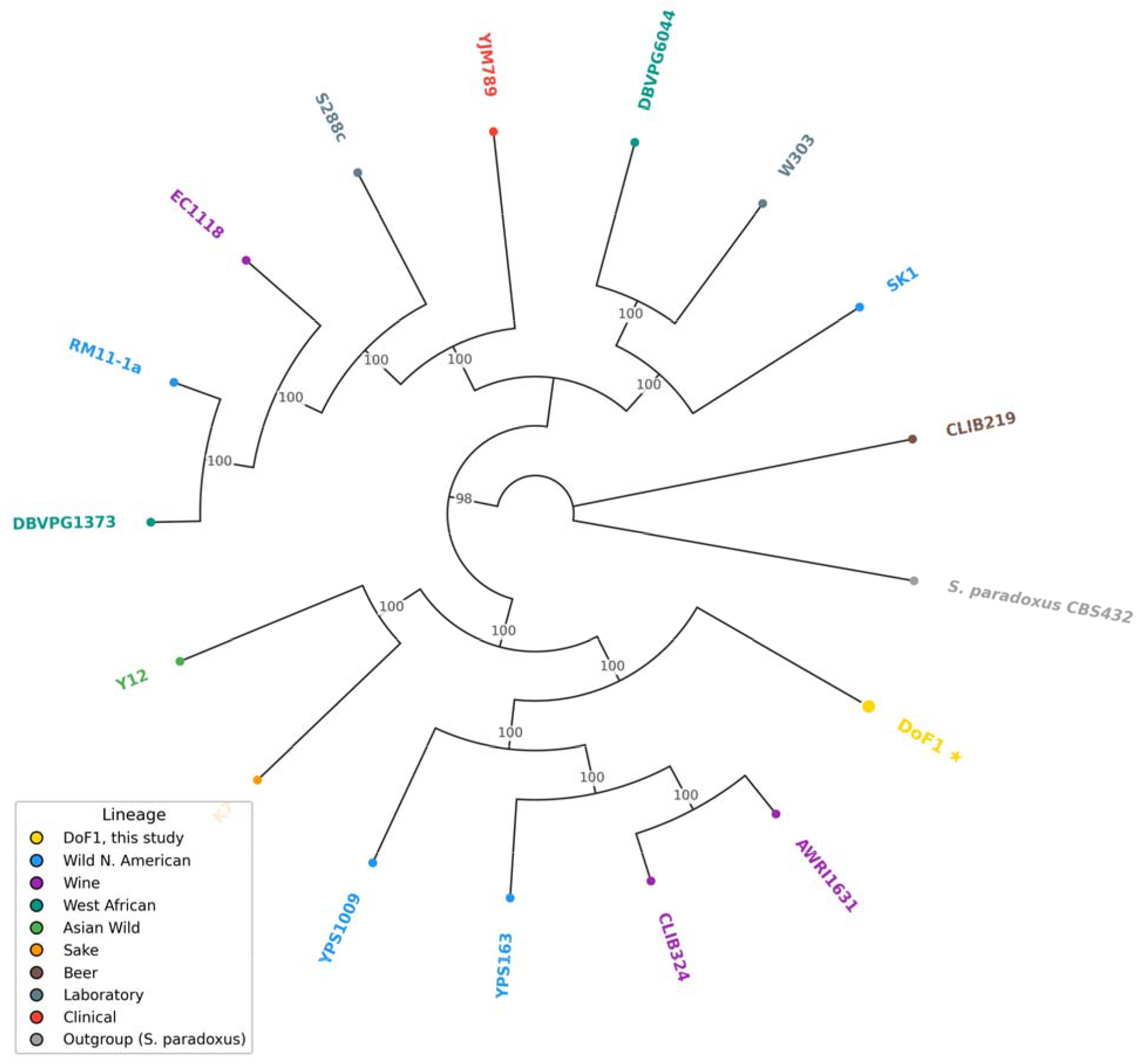
Phylogenetic placement of *S. cerevisiae* DoF1 within the broader *S. cerevisiae* species tree. Maximum likelihood circular cladogram inferred with IQ-TREE3 (LG+G4 model, 1,000 ultrafast bootstrap replicates) using 832 shared single-copy BUSCO orthologs (saccharomycetes_odb10) across 17 total taxa. Tree is rooted on *S. paradoxus* (CBS432). Bootstrap values ≥70 are shown at internal nodes. Branch lengths are equalized for display. Strain labels are colored by lineage: blue, wild North American; purple, wine; teal, West African; green, Asian wild; orange, sake; brown, beer; gray, laboratory; red, clinical; light gray, outgroup (*S. paradoxus*). DoF1 (gold star, ⍰) places within a well-supported clade (100% bootstrap) comprising wild North American strains (YPS163, YPS1009) and wine-lineage strains (AWRI1631, CLIB324), entirely separate from all laboratory, bioethanol, and West African lineages. *S. cerevisiae* clinical isolates are phylogenetically scattered across the species tree rather than forming a monophyletic lineage (21), and this placement is consistent with wild environmental provenance.

### Sensory Profile of the Declaration of Fermentation Ale

Volunteer panels at two independent public tasting events — both convened by the Indiana University Arbutus Society — evaluated the finished ale using a structured sensory scorecard (Appendix 5). Tasting 1 comprised 19 panelists and Tasting 2 comprised 14–15 panelists per attribute. To confirm that the two panels could be legitimately pooled, scores were compared between events using the Mann–Whitney U test with Benjamini–Hochberg false discovery rate correction; no attribute differed significantly between tastings (all corrected p□>□0.05), supporting pooling into a combined dataset (combined *n* = 33–34 per attribute). The beer exhibited a fruity-funky profile consistent with wild-yeast fermentation (Fig. 2): in the combined dataset, Funky/Earthy was the dominant aroma note (mean = 4.1, SD = 3.1) and Body/Mouthfeel received the highest flavor score (5.7 ± 2.4), consistent with the full-bodied character expected from a non-domesticated strain (19). Funky/Earthy aroma showed the greatest interindividual variation (SD = 3.1) and a visibly bimodal distribution, reflecting well-documented population-level differences in detection threshold for 4-ethylphenol and related phenolic compounds (20). This variance is itself pedagogically valuable: students encounter a concrete, taste-based example of biological variation in chemosensory perception, connecting the fermentation CURE directly to concepts in human genetics and sensory physiology. Individual panelist scores from both tasting events, displayed as raincloud plots, are provided in Supplemental Figure S1.

**Figure 2.**
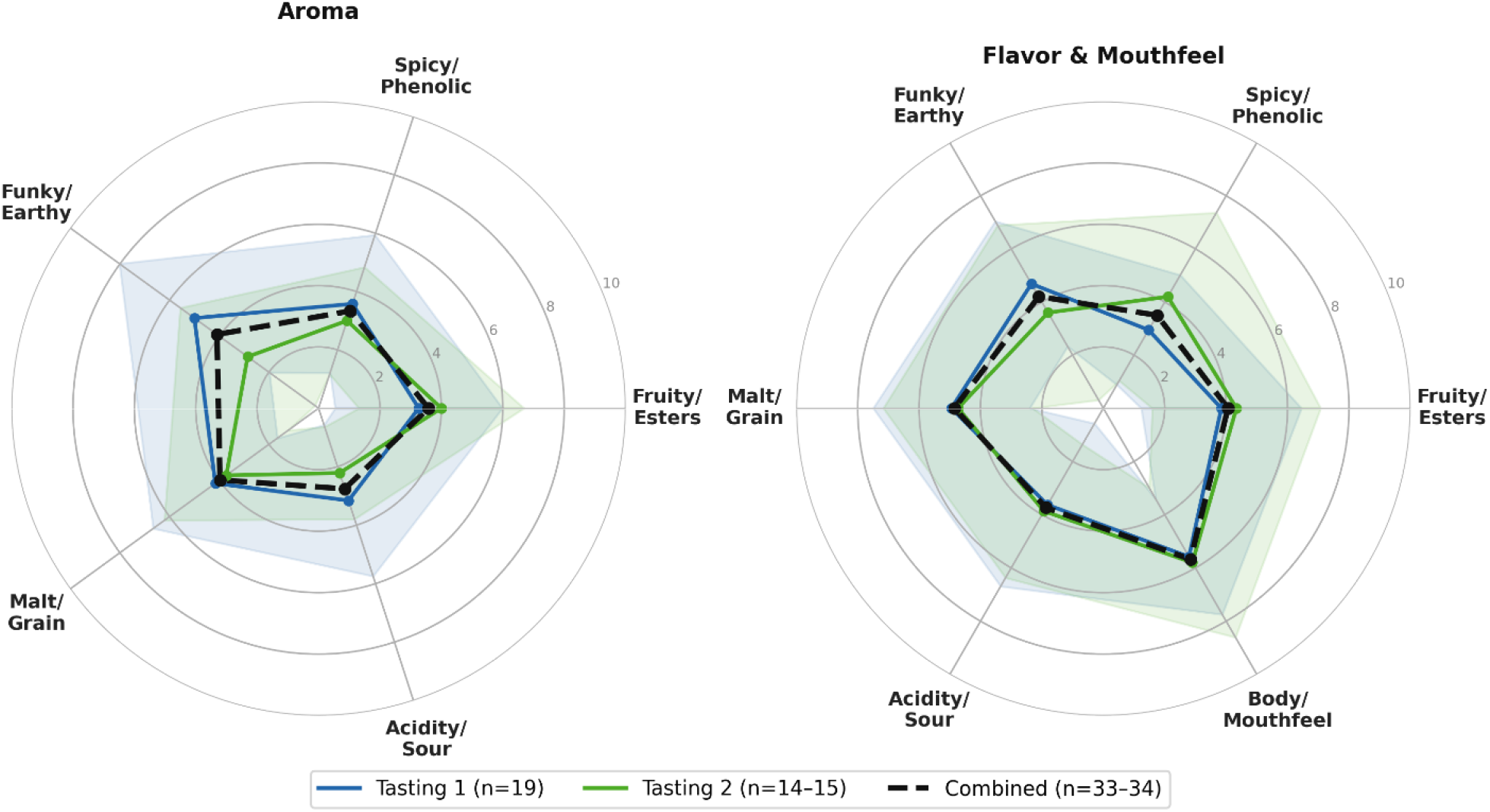
Sensory profile of the Declaration of Fermentation Ale evaluated by a volunteer tasting panel. Radar charts display mean attribute scores (bold line) with ±1 SD (shading) for Aroma (left) and Flavor & Mouthfeel (right) on a 0–10 intensity scale (0_□=_□none, 10_□=_□high). Blue = Tasting 1 (n_□=_□19); green = Tasting 2 (*n* = 18–19) and Flavor & Mouthfeel (right; *n* = 19–20). Attributes were rated on a 0–10 intensity scale (0 = none, 10 = high). Individual panelist scores are provided in Supplemental Figure S1.

## DISCUSSION AND NEXT STEPS

The Declaration of Fermentation project demonstrates that a community-embedded, historically contextualized wild yeast CURE is scientifically feasible and logistically achievable within a standard academic calendar. The core result – that a campus-isolated wild *S. cerevisiae* strain, confirmed by WGS as a genuine environmental isolate, can produce a publicly available ale with a sensory profile characteristic of wild-strain fermentation – validates the scientific premise of the curriculum. The fruity-funky profile of *S. cerevisiae* DoF1, with Funky/Earthy as the dominant aroma note and a full-bodied mouthfeel, is directly distinguishable from the clean, neutral profile of domesticated laboratory strains (19). This is a distinction students can taste, which makes the concept of phenotypic diversity in microorganisms viscerally concrete rather than abstractly described. Phylogenetically, DoF1 places within a well-supported clade (100% bootstrap) comprising wild North American strains and wine-lineage strains, separate from all laboratory, bioethanol, and West African lineages in the reference panel (Fig. 1). *S. cerevisiae* clinical isolates are phylogenetically scattered across the species tree rather than forming a monophyletic pathogenic lineage (21). The DoF1-YJM993 sister relationship therefore reflects shared wild-origin ancestry, not membership in a pathogenic lineage. *S. cerevisiae* is not considered a food safety risk in healthy individuals (18), and the wild-isolate status of DoF1 is fully consistent with its use in a public fermentation product.

This project builds explicitly on the two prior wild yeast brewing CUREs. Scholes *et al*. (2021) and DeHaven *et al*. (2022) established the educational viability of wild yeast isolation and fermentation as a CURE platform and reported quantitative learning gains (10, 11). Our contribution is not to replicate those findings but to extend the model in three directions. First, the inclusion of WGS-confirmed strain identity elevates the scientific standard: DoF1 is not merely “a wild yeast” but a characterized environmental isolate that can be deposited, cited, and used by others. This transforms the strain from a pedagogical tool into a scientific object, giving the CURE genuine novelty value in both educational and research contexts. Second, the brewery partnership structures community engagement as a curricular requirement rather than an optional showcase, which changes the epistemic stakes for students. Their isolate will be used by professionals, evaluated by the public, and consumed at a civic event. Third, the semi-quincentennial framing provided a non-arbitrary reason for the project to exist at this specific time and place, connecting microbiology to history, civic identity, and community celebration in a way that a generic “brew something with your isolate” framework cannot.

The high interindividual variance in Funky/Earthy scores is itself a pedagogically valuable result that neither prior wild yeast CURE (10, 11) was positioned to observe, because neither included a structured public sensory evaluation. Replication across two independent tasting events — with no statistically significant difference on any of 11 scored attributes (Mann–Whitney U, Benjamini–Hochberg-corrected p□>□0.05) — further validates the scorecard instrument and confirms consistency of the beer across service occasions. The bimodal distribution of 4-ethylphenol sensitivity across the combined panel (n□=□33–34) provides instructors with a concrete, student-generated example of biological variation in chemosensory perception, directly connecting the fermentation CURE to broader concepts in human genetics and sensory physiology (20). We suggest that structured community sensory evaluation – using a validated scorecard instrument conducted with a naive panel at multiple events – is a scalable addition to any wild yeast CURE that adds scientific content, community engagement, and pedagogical richness at minimal additional cost.

The modular design of the curriculum allows the depth and technical complexity to be adjusted for different course levels and resource environments. At the core, the CURE requires only standard microbiological equipment: enrichment medium, solid growth medium, incubator, microscope, and small fermentation vessels. Molecular identification is optional but strongly recommended, as species-level confirmation is what distinguishes authentic bioprospecting from a qualitative observation exercise. WGS is an advanced extension appropriate for courses with access to genomics resources or funds for external sequencing services (*e*.*g*., Plasmidsaurus; Eugene, OR). The brewery partnership is the most logistically demanding element, but analogous community partnerships – with campus dining services, local food producers, or public health laboratories – could provide equivalent community embedding without requiring a licensed brewing operation.

Perhaps the most consequential scalability feature of this framework is its natural amenability to interdisciplinary implementation. The Declaration of Fermentation project, as piloted, was executed by microbiologists and molecular biologists, but the full arc of the CURE maps cleanly onto the expertise of students across a university. History students could identify and translate period-appropriate recipes from primary sources, contextualizing the fermentation science within material culture and foodways scholarship. Food science students could contribute to recipe formulation, ingredient sourcing, and quality analysis of the finished product. Field biology or ecology students could lead or co-lead the environmental sampling component, applying community ecology principles to microhabitat selection and sampling design. Microbiology and molecular biology students would handle enrichment, isolation, and rDNA-based identification. Students in bioinformatics or computational biology courses could take the sequencing data downstream, performing phylogenetic placement, variant calling, or comparative genomic analysis. If a finished product is produced, business, marketing, communications, and event management students could design and execute a launch campaign, manage stakeholder relationships with the brewery partner, and evaluate community impact. In a fully realized interdisciplinary implementation, the proof-of-principle limitation we acknowledge below – that graduate researchers rather than undergraduates piloted the wet lab work – dissolves: the scientific validation is distributed across concurrent courses, each contributing its disciplinary expertise to a shared research object. This model aligns with emerging frameworks for cross-departmental CUREs (22, 23) and represents a natural extension of the place-based, community-embedded design principles that distinguish this project from prior wild yeast brewing curricula (10, 11).

The place-based narrative principle generalizes beyond the specific historical moment of the American semi-quincentennial and beyond beer as the fermentation vehicle. For instance, institutions near sites associated with Johnny Appleseed could build an equivalent CURE around wild yeast isolation from heritage apple orchards, producing a hard cider with documented regional provenance. Institutions in the mid-Atlantic or Appalachian regions could connect to colonial honey production and produce mead; those in the Pacific Northwest could partner with Indigenous food sovereignty initiatives around traditional fermented foods. What the Declaration of Fermentation project demonstrates is not that colonial ale is the right product; it is that a locally meaningful fermentation narrative, a genomically characterized wild isolate, and a community partnership that puts student research in front of a public audience are design principles that travel. The specific history, organism, and product are variables; the framework is the contribution.

### Limitations

Several important limitations of this proof-of-concept implementation should be acknowledged. The full CURE framework as described here was piloted by graduate researchers rather than undergraduate students, so we cannot yet report formal learning outcome data, success rates of isolation by novice microbiologists, or timeline feasibility for students with limited laboratory experience. The core bioprospecting methodology has, however, been implemented successfully with undergraduates in prior work from this laboratory (5, 6, 9), supporting the feasibility of the individual modules even in the absence of CURE-specific assessment data. Sensory evaluation was conducted with convenience samples of event volunteers at two independent tasting events (combined n = 33–34 per attribute) rather than a trained or pre-screened panel. Scores did not differ significantly between events (Mann–Whitney U, Benjamini–Hochberg-corrected p□>□0.05 for all attributes), supporting the reliability of the instrument in community-engagement contexts, though the panel composition and setting limit generalizability to trained sensory evaluation. The genomic characterization of *S. cerevisiae* DoF1 is based on sequencing data currently under review; the strain’s phylogenetic placement among wild *vs*. domesticated *S. cerevisiae* lineages should be confirmed by deposition and independent verification before strong provenance claims are made in subsequent publications. The reference panel used for phylogenetic placement (16 *S. cerevisiae* strains) captures the major named lineages but remains small relative to the 1,011-genome dataset of Peter *et al*. (21), and placement within the current panel is well-supported but could be refined with broader taxon sampling. Expanded phylogenetic analysis is planned as part of a companion genomics study. Finally, formal undergraduate implementation, IRB-approved learning outcome assessment, and comparison with the Scholes *et al*. (10) and DeHaven *et al*. (11) frameworks are all planned for future work and are prerequisites for claims about the educational efficacy of this specific model relative to existing alternatives.

## ACKNOWLEDGMENTS

We thank Clay Steenbergen (Upland Brewing Company), Matt Wisley (Bloomington Brewing Company), Dan Dutcher (Heartwork Brewing), Chris Paumi (The Tap), Kim Collins (Towaki Brewing Company) and the rest of the Bloomington, Indiana craft beer professionals for brewing the Declaration of Fermentation. We also thank Brian Yeley and the Indiana University Foundation for organizing tasting events.

## SUPPLEMENTAL MATERIALS

**Supplemental Figure S1**. Individual panelist sensory scores for the Declaration of Fermentation Ale from two independent tasting events.

**Supplemental Figure S2**. Overview of the yeast isolation and molecular identification workflow for the Declaration of Fermentation CURE.

**Appendix 1**. Environmental sampling and enrichment protocol.

**Appendix 2**. Yeast isolation, purification, and colony characterization protocol.

**Appendix 3**. Molecular identification protocol (D1/D2 rDNA PCR and Sanger sequencing).

**Appendix 4**. Suggested assessment rubrics and learning objective mapping.

**Appendix 5**. Wild Yeast Sensory Evaluation Scorecard.

**Appendix 6**. Interdisciplinary implementation model for the Declaration of Fermentation CURE framework.

## Notes

**Support**: This work was funded by the College of Arts and Sciences, Indiana University-Bloomington and the Walter Center for Career Achievement.

**Conflict of interest**: The authors declare no conflict of interest.

https://doi.org/10.5281/zenodo.20679384

### Competing Interest Statement

The authors have declared no competing interest.

## REFERENCES

1. Auchincloss LC, Laursen SL, Branchaw JL, Eagan K, Graham M, Hanauer DI, Lawrie G, McLinn CM, Pelaez N, Rowland S, Towns M, Trautmann NM, Varma-Nelson P, Weston TJ, Dolan EL. 2014. Assessment of course-based undergraduate research experiences: a meeting report. CBE Life Sci Educ 13:29–40.

2. Brownell SE, Kloser MJ. 2015. Toward a conceptual framework for measuring the effectiveness of course-based undergraduate research experiences in undergraduate biology. Studies in Higher Education 40:525–544.

3. Boundy-Mills K. 2012. Yeast culture collections of the world: meeting the needs of industrial researchers. J Ind Microbiol Biotechnol 39:673–80.

4. Goddard MR, Greig D. 2015. Saccharomyces cerevisiae: a nomadic yeast with no niche? FEMS Yeast Res 15.

5. Osburn K, Amaral J, Metcalf SR, Nickens DM, Rogers CM, Sausen C, Caputo R, Miller J, Li H, Tennessen JM, Bochman ML. 2018. Primary souring: A novel bacteria-free method for sour beer production. Food Microbiol 70:76–84.

6. Barry JP, Metz MS, Hughey J, Quirk A, Bochman ML. 2018. Two Novel Strains of Isolated from the Honey Bee Microbiome and Their Use in Honey Fermentation. Fermentation-Basel 4.

7. Alsammar H, Delneri D. 2020. An update on the diversity, ecology and biogeography of the Saccharomyces genus. FEMS Yeast Res 20.

8. Steensels J, Verstrepen KJ. 2014. Taming wild yeast: potential of conventional and nonconventional yeasts in industrial fermentations. Annu Rev Microbiol 68:61–80.

9. Araujo Piraine RE, Nickens DG, Sun DJ, Leivas Leite FP, Bochman ML. 2022. Isolation of wild yeasts from Olympic National Park and Moniliella megachiliensis ONP131 physiological characterization for beer fermentation. Food Microbiol 104:103974.

10. Scholes AN, Pollock ED, Lewis JA. 2021. A Wild Yeast Laboratory Activity: From Isolation to Brewing. J Microbiol Biol Educ 22.

11. DeHaven B, Sato B, Mello J, Hill T, Syed J, Patel R. 2022. Bootleg Biology: a Semester-Long CURE Using Wild Yeast to Brew Beer. J Microbiol Biol Educ 23.

12. Johnson MD, Margell ST, Goldenberg K, Palomera R, Sprowles AE. 2023. Impact of a First-Year Place-Based Learning Community on STEM Students’ Academic Achievement in their Second, Third, and Fourth Years. Innov High Educ 48:169–195.

13. Bochman ML. 2025. DoF1 phylogenomics pipeline (Version 1.0.0) Zenodo, 10.5281/zenodo.20679384.

14. Kurtzman CP, Robnett CJ. 1997. Identification of clinically important ascomycetous yeasts based on nucleotide divergence in the 5’ end of the large-subunit (26S) ribosomal DNA gene. J Clin Microbiol 35:1216–23.

15. Minh BQ, Schmidt HA, Chernomor O, Schrempf D, Woodhams MD, von Haeseler A, Lanfear R. 2020. Corrigendum to: IQ-TREE 2: New Models and Efficient Methods for Phylogenetic Inference in the Genomic Era. Mol Biol Evol 37:2461.

16. Minh BQ, Schmidt HA, Chernomor O, Schrempf D, Woodhams MD, von Haeseler A, Lanfear R. 2020. IQ-TREE 2: New Models and Efficient Methods for Phylogenetic Inference in the Genomic Era. Mol Biol Evol 37:1530–1534.

17. Manni M, Berkeley MR, Seppey M, Simao FA, Zdobnov EM. 2021. BUSCO Update: Novel and Streamlined Workflows along with Broader and Deeper Phylogenetic Coverage for Scoring of Eukaryotic, Prokaryotic, and Viral Genomes. Mol Biol Evol 38:4647–4654.

18. Sanders ME, Akkermans LM, Haller D, Hammerman C, Heimbach J, Hormannsperger G, Huys G, Levy DD, Lutgendorff F, Mack D, Phothirath P, Solano-Aguilar G, Vaughan E. 2010. Safety assessment of probiotics for human use. Gut Microbes 1:164–85.

19. Hyma KE, Saerens SM, Verstrepen KJ, Fay JC. 2011. Divergence in wine characteristics produced by wild and domesticated strains of Saccharomyces cerevisiae. FEMS Yeast Res 11:540–51.

20. Tempere S, Cuzange E, Malak J, Bougeant JC, de Revel G, Sicard G. 2011. The Training Level of Experts Influences their Detection Thresholds for Key Wine Compounds. Chemosensory Perception 4:99–115.

21. Peter J, De Chiara M, Friedrich A, Yue JX, Pflieger D, Bergstrom A, Sigwalt A, Barre B, Freel K, Llored A, Cruaud C, Labadie K, Aury JM, Istace B, Lebrigand K, Barbry P, Engelen S, Lemainque A, Wincker P, Liti G, Schacherer J. 2018. Genome evolution across 1,011 Saccharomyces cerevisiae isolates. Nature 556:339–344.

22. Shortlidge EE, Brownell SE. 2016. How to Assess Your CURE: A Practical Guide for Instructors of Course-Based Undergraduate Research Experiences. J Microbiol Biol Educ 17:399–408.

23. Dolan EL. 2016. Course-based undergraduate research experiences: current knowledge and future directions. National Academies of Sciences, Engineering, and Medicine, Washington, DC.

