## Supplementary materials for "Declaration of Fermentation: Community-Embedded Wild Yeast Bioprospecting as a Model for Place-Based CURE Design"

This document contains the following supplemental materials:

- **Figure S1.** Individual panelist sensory scores for the Declaration of Fermentation Ale (two tasting events; raincloud plots).
- **Figure S2.** Overview of the yeast isolation and molecular identification workflow for the Declaration of Fermentation CURE.
- **Appendix 1.** Environmental Sampling and Yeast Enrichment Protocol
- **Appendix 2.** Yeast Isolation, Purification, and Phenotypic Screening Protocol
- **Appendix 3.** Molecular Identification Protocol (rDNA D1/D2 PCR and Sanger Sequencing)
- **Appendix 4.** Assessment Instruments and Interdisciplinary Coordination Timeline
- **Appendix 5.** Wild Yeast Sensory Evaluation Scorecard
- **Appendix 6.** Interdisciplinary implementation model
- **Supplemental References**

**Supplemental figures**

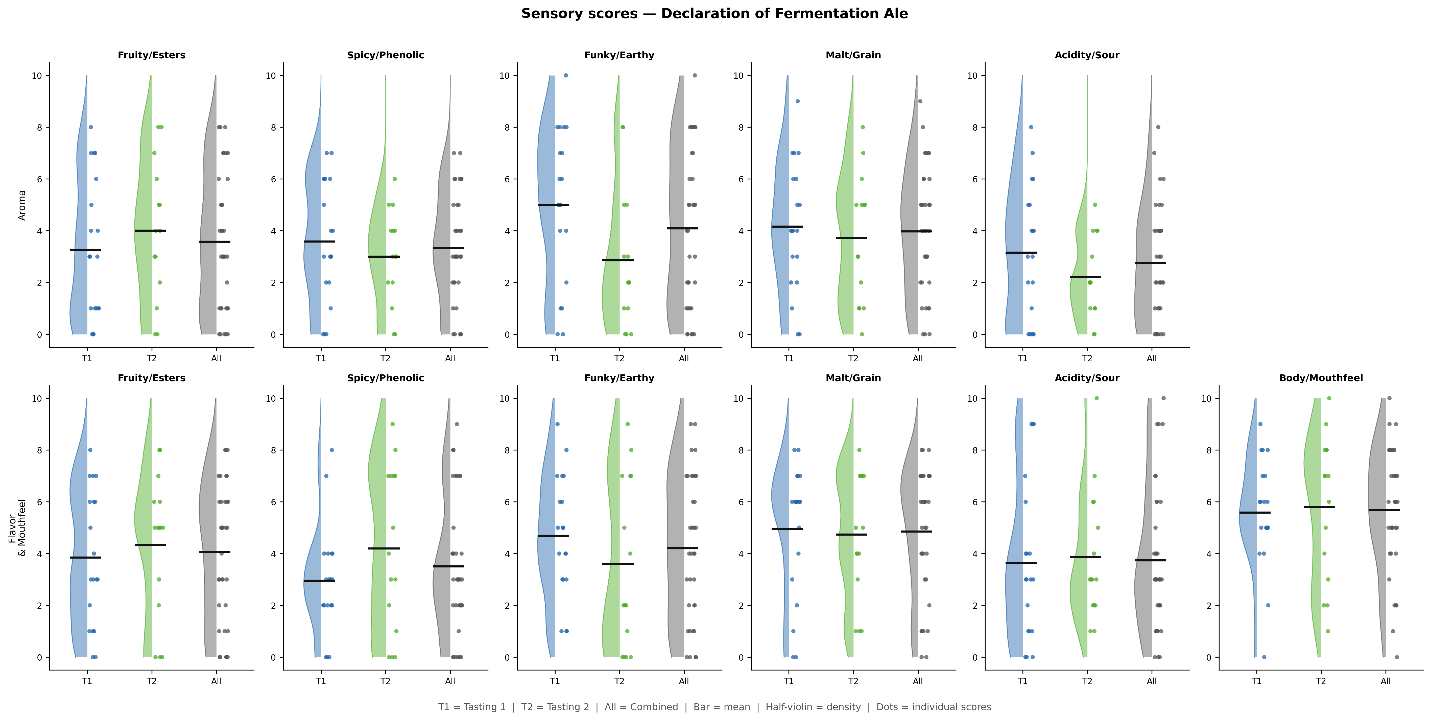

**Supplemental Figure S1.** Individual panelist sensory scores for the Declaration of Fermentation Ale from two independent tasting events. Raincloud plots show the kernel density estimate (half-violin, left), individual panelist scores (dots, right, jittered horizontally for visibility), and mean (horizontal bar) for each attribute, grouped by Tasting 1 (blue; n = 19), Tasting 2 (green; n = 14–15), and combined (gray; n = 33–34). All attributes scored on a 0–10 intensity scale (0 = none, 10 = high). No attribute differed significantly between tasting events (Mann–Whitney U test with Benjamini–Hochberg false discovery rate correction; corrected p > 0.05 for all 11 comparisons).

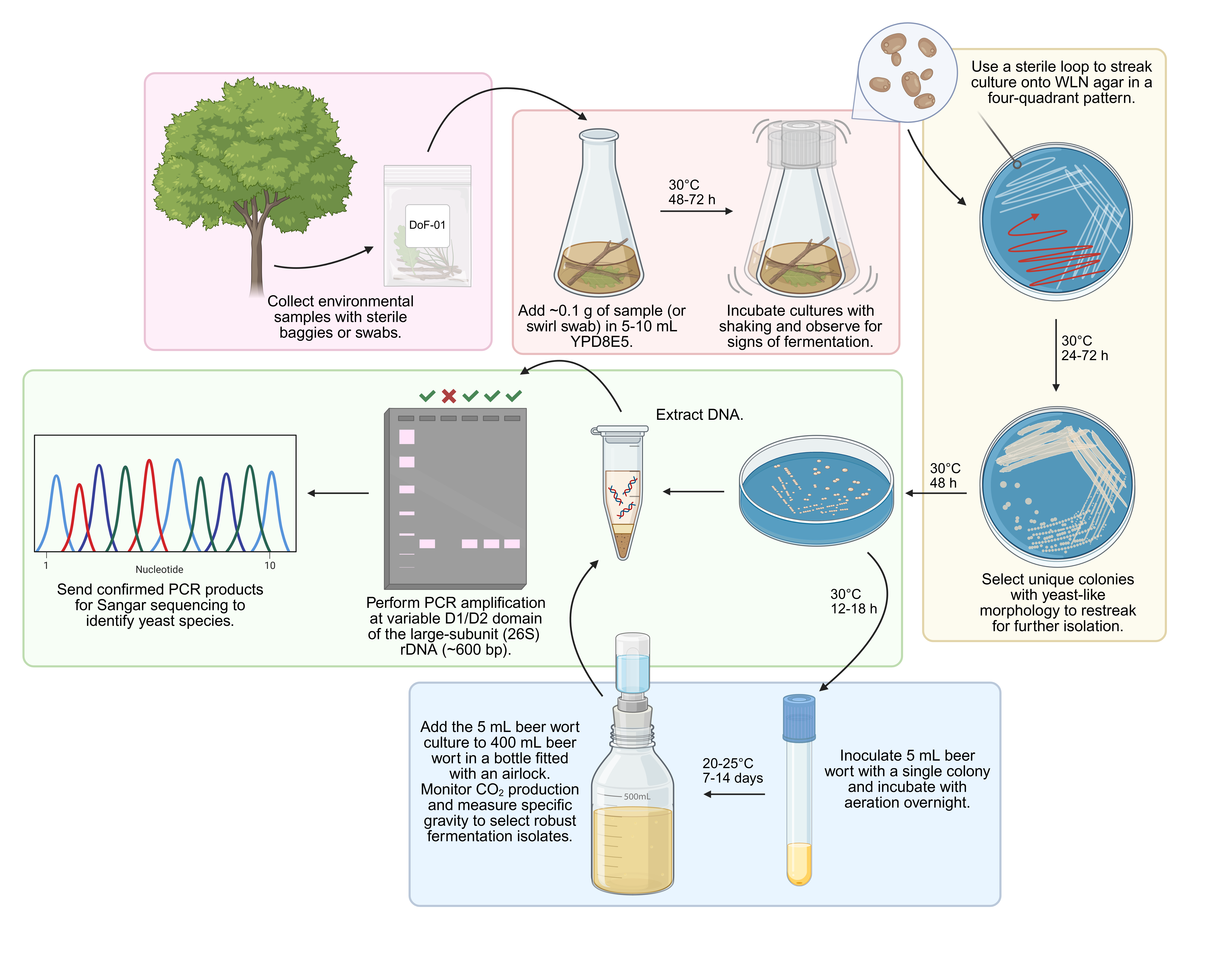

**Supplemental Figure S2.** Overview of the yeast isolation and molecular identification workflow for the Declaration of Fermentation CURE. Colored boxes indicate workflow phases: pink, environmental sampling and enrichment (Module 2); yellow, isolation and purification (Module 2); green, molecular identification by rDNA PCR and Sanger sequencing (Module 3a); blue, small-scale fermentation screening (Module 4). Timing annotations indicate approximate incubation durations at each step.

**Appendix 1: Environmental Sampling and Yeast Enrichment Protocol**

**Overview**

This protocol describes how to collect environmental samples from field sites and enrich for ethanol-tolerant yeasts using selective liquid medium. It corresponds to Module 2 of the Declaration of Fermentation CURE (Table 1 of the main text). Estimated time: 1 laboratory session (2–3 h) for sample collection and medium inoculation, followed by 48–72 h incubation.

**Learning Objectives**

Upon completing this module, students will be able to:

- Apply principles of microbial ecology to select informative environmental sampling sites.
- Collect environmental samples aseptically using appropriate materials.
- Prepare and inoculate selective enrichment medium for ethanol-tolerant yeasts.
- Maintain accurate field and laboratory records.

**Materials**

***For environmental sampling***

- New, unopened zip-type plastic bags (quart size), 2–3 per sampling site
- Sterile cotton-tipped swabs (for surface sampling)
- Permanent marker and waterproof labels
- Field data sheet (see below)
- GPS-enabled device or smartphone for coordinate recording
- Gloves (nitrile)
- Camera or smartphone for site documentation

***For enrichment medium (YPD8E5)***

Prepare the following per 1 L of enrichment medium:

- Yeast extract: 10 g
- Peptone: 20 g
- Dextrose: 80 g
- Deionized water: to 950 mL
- Autoclave at 121°C for 30 min; cool to room temperature
- After autoclaving, aseptically add: 50 mL of 100% ethanol (to reach 5% v/v final concentration)
- Antibiotics (added after autoclaving): kanamycin to a final concentration of 50 µg/mL and chloramphenicol to a final concentration of 34 µg/mL (from stock solutions of 50 mg/mL and 34 mg/mL in water and ethanol, respectively)

**[NOTE TO INSTRUCTORS:** YPD8E5 = YPD with 8% dextrose and 5% ethanol. The elevated dextrose and ethanol concentrations provide selective pressure for ethanol-tolerant yeasts while suppressing most bacteria and non-fermentative organisms.**]**

**Field Sampling Procedure**

**Step 1.** Before departing, assign a unique sample code to each planned site (*e.g*., DoF-01, DoF-02). Record GPS coordinates, date, time, weather conditions, and a brief habitat description on the field data sheet for each site.

**Step 2.** At each site, collect samples aseptically. For surfaces (*e.g*., tree bark): moisten a sterile swab with sterile water, swab a ~5 cm² area vigorously, and transfer the swab head into a labeled zip bag containing 2 mL of sterile water. For soil or leaf litter: collect approximately 1 g of material directly into a labeled zip bag using a sterile implement. Alternatively, turn the bag inside out on your hand, grab a sample, turn the carefully turn the bag right-side out, and seal.

**Step 3.** Collect 2–3 samples per site from slightly different microhabitats (*e.g*., lower *vs.* upper bark, sun-exposed *vs*. shaded surfaces). Document each with a photograph.

**Step 4.** Keep samples cool and transport to the laboratory within 4 h. Process immediately or store at 4°C overnight (no longer than 24 h).

**Enrichment Procedure**

**Step 1.** Label one test tube or flask per sample.

**Step 2.** Add 5–10 mL of YPD8E5 to each.

**Step 3.** Transfer sample material: for swab samples, agitate the swab head in the medium and squeeze against the tube wall before discarding; for soil/leaf samples, add approximately 0.1 g of material and vortex briefly.

**Step 4.** Incubate cultures at 30°C with shaking (200 rpm) for 48–72 h. Observe daily for signs of fermentation (turbidity, CO₂ bubbles, yeast-like odor).

**Step 5 (optional).** After incubation, examine a small aliquot under the microscope (40× objective) to confirm the presence of budding yeast cells before proceeding to Appendix 2. Yeast Isolation, Purification, and Phenotypic Screening Protocol.

**Field Data Sheet**

| **Sample Code** | **Date / Time** | **GPS Coordinates** | **Site Description** | **Sample Type (bark / soil / fruit / other)** | **Notes** |
| --- | --- | --- | --- | --- | --- |

**Safety**

- Work outdoors in pairs; notify a supervisor of sampling locations before departing.
- Wear gloves when handling environmental samples.
- Ethanol is flammable; add to media away from open flames.
- Dispose of enrichment cultures by autoclaving or treating with 10% bleach before discarding.

**Appendix 2: Yeast Isolation, Purification, and Phenotypic Screening Protocol**

**Overview**

This protocol describes how to isolate individual yeast colonies from enrichment cultures, purify strains, and screen them for fermentation performance. It corresponds to Modules 2–3 of the Declaration of Fermentation CURE. Estimated time: 2–3 laboratory sessions over 1–2 weeks.

**Learning Objectives**

- Perform streak-plate isolation to obtain single colonies from mixed cultures.
- Distinguish yeast colony morphologies macroscopically and microscopically.
- Screen isolates for fermentation activity using a rapid small-scale assay.
- Maintain a curated strain library with annotated records.

**Materials**

- WLN (Wallerstein Laboratory Nutrient) agar plates
- YPD agar plates (for maintenance)
- YPD liquid medium
- Sterile inoculating loops and spreaders
- Glass microscopy slides and coverslips
- Gram stain reagents (crystal violet, Gram's iodine, decolorizer, safranin)
- Methylene blue (for viability staining, optional)
- Brightfield microscope with 40× and 100× objectives
- 10-mL conical tubes, glass fermentation tubes with airlocks, or 500-mL glass bottles fitted with airlocks
- Commercial beer wort (original gravity ~1.040–1.050) or prepared wort
- *Note:* WLN agar produces differential colony pigmentation that aids yeast/non-yeast discrimination; green colonies are typical of *Saccharomyces* spp.

**Streak-Plate Isolation**

**Step 1.** After 48–72 h incubation of enrichment cultures, streak a loopful of each culture onto a WLN agar plate using a standard four-quadrant streak pattern.

**Step 2.** Incubate plates inverted at 30°C for 24–72 h.

**Step 3.** Examine plates and select colonies with yeast-like morphology (smooth, moist, cream to green pigmentation on WLN). For each culture, select 3–5 distinct colony morphotypes.

**Step 4.** Restreak each selected colony onto a fresh WLN or YPD agar plate to ensure purity. Incubate at 30°C for 48 h.

**Step 5.** Confirm purity by microscopy: prepare a wet mount, examine under 40× for uniform cell morphology, and under 100× (oil immersion) for budding yeast morphology. Note cell size, shape, budding pattern, and any pseudohyphae.

**Colony Documentation**

Record the following for each isolate in your laboratory notebook:

- Colony color (on WLN and YPD)
- Colony size (mm diameter after 24 and 48 h)
- Colony texture (smooth, rough, wrinkled, mucoid)
- Colony margin (entire, undulate, filamentous)
- Microscopic morphology (cell shape, size, budding pattern)
- Gram reaction (yeast cells should appear Gram-positive, purple)

**Small-Scale Fermentation Screening**

**Step 1.** Inoculate each purified isolate into 5 mL of beer wort in a capped tube with a small airlock or cotton plug. Alternatively, add a 5-mL overnight culture grown at 30°C with aeration in wort to 400 mL wort in a 500-mL bottle topped with a rubber stopper fitted with a homebrew-style airlock.

**Step 2.** Incubate at 20–25°C for 7-14 days (ale fermentation temperatures).

**Step 3.** Monitor CO₂ production daily by observing airlock activity. Record qualitative fermentation intensity (none / low / moderate / high).

**Step 4.** On days 1, 3, 5, and 7, measure specific gravity using a digital refractometer and record values. Calculate apparent attenuation: [(OG − FG) / (OG − 1)] × 100, where OG = original gravity and FG = final gravity.

**Step 5.** Select isolates with robust fermentation performance (high attenuation, active CO₂ production) for molecular identification (Supplemental Material S3).

**Strain Library**

| **Isolate ID** | **Source Sample** | **Colony Morphology (WLN)** | **Gram Stain** | **Microscopy Notes** | **Fermentation Score (none/low/mod/high)** | **Selected for ID?** |
| --- | --- | --- | --- | --- | --- | --- |

**Appendix 3: Molecular Identification Protocol (rDNA D1/D2 PCR and Sanger Sequencing)**

**Overview**

This protocol describes DNA extraction from yeast cultures and PCR amplification of the variable D1/D2 domain of the large-subunit (26S) rDNA for Sanger sequencing and BLASTN-based species identification. This approach provides reliable species-level identification for most yeast genera (1, 2). It corresponds to Module 3a of the Declaration of Fermentation CURE.

**Learning Objectives**

- Extract genomic DNA from yeast using a standard lysis protocol.
- Design and interpret PCR amplification using universal yeast primers NL1 and NL4.
- Assess PCR product quality by agarose gel electrophoresis.
- Submit samples for Sanger sequencing and interpret BLASTN results.
- Assign species identity using percent sequence identity criteria.

**Materials**

***DNA extraction***

- Commercial yeast DNA extraction kit (e.g., YeaStar Genomic DNA Kit; Zymo Research)
  - Alternatively: 200 µL of 0.2 M lithium acetate/1% SDS, 65°C water bath (or thermomixer or heat block), and 100% isopropanol
- Microcentrifuge tubes (1.5 mL)
- Microcentrifuge

***PCR amplification***

- Universal primers: NL1 (5ʹ-GCATATCAATAAGCGGAGGAAAAG-3ʹ) and NL4 (5ʹ-GGTCCGTGTTTCAAGACGG-3ʹ)
- PCR master mix (*Taq* polymerase, dNTPs, MgCl₂, reaction buffer)
- PCR tubes or strip tubes
- Thermal cycler

***Gel electrophoresis and sequencing***

- 1% agarose gel in 1× TAE or TBE buffer
- SYBR Safe or ethidium bromide for DNA visualization
- 100-bp DNA ladder
- PCR purification kit (QIAquick PCR Purification Kit; Qiagen)
- Sanger sequencing service (Eurofins Genomics)

**DNA Extraction Procedure**

***Rapid colony PCR method (recommended for screening)***

**Step 1.** Add 100 µL YPD overnight culture to 200 uL of 0.2 M lithium acetate/1% SDS in a 1.5-mL microcentrifuge tube.

**Step 2.** Incubate at 65°C for 15-20 min, then add 1000 µL of 100% isopropanol. Vortex briefly.

**Step 3.** Centrifuge at 13,000 × g for 5 min at room temperature. Decant the isopropanol, and remove remaining traces by pipetting.

**Step 4.** Quickly resuspend the pellet in 100 µL deionized water by pipetting and vortexing. Pellet cell debris as above.

**Step 5.** Transfer 1 µL of the supernatant to a fresh tube for use as PCR template. (Do not transfer the pellet.)

*Alternatively:* use a commercial yeast DNA extraction kit according to the manufacturer's instructions. Elute in 50–100 µL of elution buffer. Use 1–2 µL per PCR reaction.

**PCR Amplification**

Prepare the following reaction mix per sample:

| **Component** | **Volume (µL)** | **Final Concentration** |
| --- | --- | --- |
| 2× PCR Master Mix | 12.5 | 1× |
| NL1 primer (10 µM) | 1.0 | 0.4 µM |
| NL4 primer (10 µM) | 1.0 | 0.4 µM |
| DNA template | 1.0 | variable |
| Nuclease-free water | 9.5 | — |
| Total | 25.0 |  |

Thermal cycler conditions:

| **Step** | **Temperature** | **Time** | **Cycles** |
| --- | --- | --- | --- |
| Initial denaturation | 98°C | 5 min | 1 |
| Denaturation | 98°C | 30 s | 35 |
| Annealing | 55°C | 30 s |  |
| Extension | 72°C | 60 s |  |
| Final extension | 72°C | 5 min | 1 |
| Hold | 4°C | ∞ |  |

Expected product size: ~600 bp. Always include a negative control (no template) and a positive control (known *S. cerevisiae* DNA).

**Gel Electrophoresis and Purification**

**Step 1.** Load 5 µL of each PCR product alongside a 100 bp DNA ladder on a 1% agarose gel. Run at 100 V for 30–40 min.

**Step 2.** Visualize and photograph the gel under UV or blue light. A single band at ~600 bp indicates successful amplification.

**Step 3.** Purify successful PCR products using a commercial PCR cleanup kit according to the manufacturer's instructions. Elute in 30-50 µL of elution buffer.

**Sanger Sequencing and Species Identification**

**Step 1.** Submit purified PCR products for Sanger sequencing using primer NL1. Follow the submission instructions for your chosen sequencing provider.

**Step 2.** Retrieve sequencing results (typically as a .ab1 or .txt file). Trim low-quality ends using chromatogram inspection software (*e.g.*, FinchTV, Sequence Scanner).

**Step 3.** Navigate to NCBI BLAST (https://blast.ncbi.nlm.nih.gov/). Select BLASTN. Paste the trimmed sequence into the query box. Select "Nucleotide collection (nr/nt)" as the database and click BLAST.

**Step 4.** Examine the top hits. Record the species name, percent identity, query coverage, and accession number for the top three hits. Species identity is assigned based on ≥99% sequence identity and ≥95% query coverage to a reference sequence.

*Note on interpretation:* Some environmental yeast species cannot be reliably distinguished by D1/D2 sequencing alone; where percent identity to two or more species is ≥99%, note both possibilities and consult the ITS region for additional resolution if needed.

**Appendix 4: Assessment Instruments and Interdisciplinary Coordination Timeline**

**Section A: Learning Objectives Mapping**

The following table maps each curriculum module to the CURE learning objectives and to relevant ASM Curriculum Guidelines for Undergraduate Microbiology Education competencies.

| **Module** | **Primary Learning Objectives** | **Assessment** | **Relevant ASM Competency** |
| --- | --- | --- | --- |
| 1 (Framing & Site Selection) | Connect microbiology to historical and cultural context; evaluate field sites using ecological criteria; formulate a sampling hypothesis | Written site selection rationale; annotated historical recipe | Evolution; Microbial Ecology |
| 2 (Sampling & Isolation) | Apply aseptic technique; collect and document environmental samples; isolate yeast using enrichment and streak-plate methods | Laboratory notebook; colony morphology data table; microscopy images | Cell Biology; Metabolic Pathways |
| 3a (Molecular ID) | Extract DNA; design and troubleshoot PCR; interpret sequencing results; assign species identity using BLASTN | Annotated gel image; BLASTN results report; species ID summary | Information Flow; Evolution |
| 3b (Phylogenomics; optional) | Assess genome assembly quality; run phylogenomic pipeline; interpret phylogenetic placement in population context | BUSCO report; annotated tree figure; draft methods paragraph | Evolution; Information Flow |
| 4 (Fermentation & Sensory) | Monitor fermentation by specific gravity; perform sensory evaluation; summarize data with descriptive statistics | Fermentation graph; sensory scorecards; written fermentation report | Metabolic Pathways; Cell Biology |
| 5 (Full-Scale; optional) | Communicate professionally with external partners; manage a multi-stakeholder project; conduct community sensory panel | Project management log; sensory dataset; public-facing poster | Microbial Systems; Applied Microbiology |
| 6 (Capstone) | Synthesize data from multiple modules; communicate scientific findings to diverse audiences; evaluate peer work | Poster or oral presentation; peer review; reflection essay | Scientific Process; Communication |

**Section B: Pre/Post CURE Assessment Instrument**

The following items assess student attitudes toward research and learning gains before and after the CURE. Administer at the first and last class meetings. Responses use a 5-point Likert scale (1 = Strongly Disagree, 5 = Strongly Agree) unless otherwise noted.

***Attitudes toward research (adapted from CURE Survey, Lopatto 2004)***

- I feel like a scientist when I work in the laboratory.
- I understand how scientific research works.
- I can troubleshoot problems I encounter in the laboratory.
- My research could contribute to knowledge beyond this classroom.
- I am interested in pursuing a career in science.
- Working with real organisms that produce unpredictable results is valuable.

***Conceptual knowledge items (open response or multiple choice — instructors should customize)***

- Explain why ethanol is used in YPD8E5 enrichment medium.
- What information does the D1/D2 rDNA sequence provide about a yeast isolate? What does it not tell you?
- Why might two strains of *S. cerevisiae* isolated from the same tree produce beers with different flavor profiles?
- What is the difference between a CURE and a traditional laboratory exercise?
- Describe one way that wild yeast genetics and human sensory perception are connected.

***Community engagement item (post only)***

- This project connected my laboratory work to my community. (Likert 1–5)
- I would recommend this project to other students. (Likert 1–5)
- [Open response] Describe in 2–3 sentences what this project meant to you beyond the course grade.

**Section C: Interdisciplinary Coordination Timeline**

The following timeline is for a two-semester implementation involving multiple disciplinary cohorts. Single-semester implementations should focus on Modules 1–4 and compress or eliminate Modules 5–6.

| **Milestone** | **Responsible Cohort(s)** | **Deliverable** | **Approximate Timing** | **Downstream Dependency** |
| --- | --- | --- | --- | --- |
| Kickoff meeting: all cohorts introduced to project narrative and shared goals | All cohorts + Faculty coordinators | Shared project charter document | Semester 1, Week 1 | All subsequent modules |
| Historical recipe research complete | History / American Studies | Annotated recipe + context memo | Semester 1, Week 6 | Food Science (recipe formulation, Module 4) |
| Environmental sampling complete; enrichment cultures initiated | Field Biology / Ecology + Microbiology | Sampling data sheet; labeled enrichment cultures | Semester 1, Weeks 2–4 | Microbiology (isolation, Module 2) |
| Strain isolation and rDNA identification complete | Microbiology / Molecular Biology | Strain library; BLASTN identification report; glycerol stocks | Semester 1, Weeks 4–8 | Food Science (Module 4); Bioinformatics (Module 3b) |
| Phylogenomic analysis complete (optional) | Bioinformatics / Computational Biology | Assembly report; phylogenetic tree figure; methods paragraph | Semester 1, Weeks 6–12 | Marketing narrative; manuscript |
| Small-scale fermentation and sensory evaluation complete | Food Science + Microbiology | Fermentation data; sensory scorecard dataset | Semester 1, Weeks 6–12 | Brewery transfer (Module 5); Marketing team |
| Commercial-scale brew in progress (optional) | Food Science + Brewery partner | Fermentation log; quality control data | Semester 2, Weeks 1–6 | Public tasting event (Module 5) |
| Marketing campaign designed and approved | Business / Marketing / Communications | Brand identity; event plan | Semester 2, Weeks 8–12 | Public tasting event logistics |
| Public tasting event and volunteer sensory panel | Events Management + all cohorts | Panelist scorecards; event documentation | Semester 2, Week 14 | Capstone presentations (Module 6) |
| Capstone presentations and manuscript draft | All cohorts | Poster / oral presentations; draft manuscript sections | Semester 2, Weeks 15–16 | Final assessment; potential publication |

**Appendix 5: Wild Yeast Sensory Evaluation Scorecard**

The following instrument was used to evaluate the Declaration of Fermentation Ale at the public tasting event. It may be reproduced and adapted freely for educational use. Instructors should print one scorecard per panelist and collect completed forms immediately after the tasting.

**Instructions for Panelists**

Thank you for participating in this sensory evaluation. You will evaluate one beer sample for its aromas and flavors. Please read all instructions before beginning. Rinse your mouth with water and eat a plain cracker between phases.

Sample Code: ___________________________

**PHASE 1: AROMA (Smell Only)**

Swirl the glass gently and take 2–3 short sniffs without inserting your nose into the glass. Rate the intensity of each aroma from 0 (none) to 10 (very high).

| **Attribute** | **Descriptors** | **Score (0 = None → 10 = High)** |
| --- | --- | --- |
| Fruity / Esters | Apple, pear, stone fruit, tropical notes | 0 1 2 3 4 5 6 7 8 9 10 |
| Spicy / Phenolic | Clove, pepper, medicinal, anise | 0 1 2 3 4 5 6 7 8 9 10 |
| Funky / Earthy | Barnyard, hay, leather, horse blanket | 0 1 2 3 4 5 6 7 8 9 10 |
| Malt / Grain | Sweet corn, cracker, biscuit, cereal | 0 1 2 3 4 5 6 7 8 9 10 |
| Acidity / Sour | Tartness, lactic, lemon-like | 0 1 2 3 4 5 6 7 8 9 10 |

Aroma notes: _______________________________________________________________________________

**PHASE 2: FLAVOR & MOUTHFEEL (Taste)**

Take a sip large enough to coat your tongue. Swallow (or expectorate). Rate the intensity of each attribute from 0 (none / very light) to 10 (very high / very heavy).

| **Attribute** | **Descriptors** | **Score (0 = None → 10 = High)** |
| --- | --- | --- |
| Fruity / Esters | Apple, pear, stone fruit, tropical notes | 0 1 2 3 4 5 6 7 8 9 10 |
| Spicy / Phenolic | Clove, pepper, medicinal, anise | 0 1 2 3 4 5 6 7 8 9 10 |
| Funky / Earthy | Barnyard, hay, leather, horse blanket | 0 1 2 3 4 5 6 7 8 9 10 |
| Malt / Grain | Sweet corn, cracker, biscuit, cereal | 0 1 2 3 4 5 6 7 8 9 10 |
| Acidity / Sour | Tartness, lactic, lemon-like | 0 1 2 3 4 5 6 7 8 9 10 |
| Body / Mouthfeel | Light/thin → full/heavy, slick, astringent | 0 1 2 3 4 5 6 7 8 9 10 |

Flavor / mouthfeel notes: ______________________________________________________________________

Overall impression (circle one): Did not enjoy	Neutral	Enjoyed	Really enjoyed

General comments: _____________________________________________________________________________

**Instructor note:** Please rinse your mouth with water and eat a plain cracker between samples if evaluating more than one beer. Return completed scorecard to the table facilitator.

**Appendix 6. Interdisciplinary Implementation Model**

The Declaration of Fermentation CURE maps naturally onto the expertise of students across multiple disciplines. The table below outlines representative disciplinary contributions, primary deliverables, and integration points for a full interdisciplinary implementation. Representative course names are illustrative; equivalent courses at home institutions may be substituted. Semester and week assignments reflect the full two-semester track; single-semester implementations should prioritize Modules 1–4.

| **Discipline** | **Representative Course** | **Contribution to CURE** | **Primary Deliverable** | **Integration Point** | **Semester** |
| --- | --- | --- | --- | --- | --- |
| History / American Studies | Food, Culture & History; Colonial American Life | Identify, translate, and modernize 18th-century ale recipes from primary sources; contextualize brewing within colonial foodways and material culture | Annotated historical recipe with modernization rationale; 2-page historical context memo | Recipe delivered to Food Science / Microbiology teams before fermentation planning begins (Module 1 → Module 4) | Sem. 1, Wks 1–6 |
| Field Biology / Ecology | Environmental Microbiology; Field Methods in Ecology | Lead environmental sampling using community ecology principles; select microhabitats; document sampling metadata; co-lead enrichment | Sampling design document; field data sheet with GPS coordinates, habitat notes, and sample metadata | Samples handed off to Microbiology team for enrichment and isolation (Module 2) | Sem. 1, Wks 2–4 |
| Microbiology / Molecular Biology | Microbiology Laboratory; Molecular Biology Lab | Enrichment, isolation, and colony characterization; rDNA-based species identification; strain curation and maintenance | Strain library with annotated colony characteristics; BLASTN identification report; frozen glycerol stocks | Validated strain transferred to Food Science and Bioinformatics teams (Module 3a → Modules 3b and 4) | Sem. 1, Wks 4–8 |
| Bioinformatics / Computational Biology | Bioinformatics; Genomics and Sequence Analysis | Whole-genome sequencing coordination; BUSCO assembly assessment; AUGUSTUS annotation; phylogenomic analysis; interpretation of wild vs. domesticated placement | Assembly completeness report; annotated phylogenetic tree figure; written genomics methods paragraph | Genomic confirmation of wild-isolate status informs Marketing team narrative and manuscript (Module 3b) | Sem. 1, Wks 6–12 |
| Food Science | Fermentation Science; Sensory Evaluation | Recipe formulation with brewer; small-scale and commercial fermentation monitoring; sensory scorecard design; volunteer panel coordination; data analysis | Fermentation data table and graph; validated sensory scorecard; panel dataset with means and SDs | Fermentation results and sensory data delivered to Marketing team and manuscript (Module 4 → Module 5) | Sem. 1–2, Wks 6–14 |
| Business / Marketing / Communications | Brand Management; Science Communication | Design product name and visual identity; develop public launch campaign; coordinate brewery partnership communications; assess community engagement | Brand identity package; event plan with timeline; post-event engagement report; social media content | Campaign launches at public tasting event, integrating all prior deliverables (Module 5 capstone) | Sem. 2, Wks 10–16 |
| Events Management | Event Planning and Logistics | Venue coordination; volunteer panel recruitment; scorecard distribution and collection; logistics for public celebration event | Event run-of-show document; completed panelist scorecards; post-event logistics debrief | Tasting event is the integrative capstone at which all disciplinary threads converge (Module 6) | Sem. 2, Wks 12–16 |

**SUPPLEMENTAL REFERENCES**

1.	Kurtzman CP, Robnett CJ. 1997. Identification of clinically important ascomycetous yeasts based on nucleotide divergence in the 5' end of the large-subunit (26S) ribosomal DNA gene. J Clin Microbiol 35:1216–23.

2.	Osburn K, Amaral J, Metcalf SR, Nickens DM, Rogers CM, Sausen C, Caputo R, Miller J, Li H, Tennessen JM, Bochman ML. 2018. Primary souring: A novel bacteria-free method for sour beer production. Food Microbiol 70:76–84.
